# Efficient computation of co-transcriptional RNA-ligand interaction dynamics

**DOI:** 10.1101/248450

**Authors:** Michael T. Wolfinger, Christoph Flamm, Ivo L. Hofaeker

**Author notes:** Email addresses:* (Michael T. Wolfinger), (Christoph Flamm), (Ivo L. Hofacker).

## Abstract

Riboswitches form an abundant class of cis-regulatory RNA elements that mediate gene expression by binding a small metabolite. For synthetic biology applications, they are becoming cheap and accessible systems for selectively triggering transcription or translation of downstream genes. Many riboswitches are kinetically controlled, hence knowledge of their co-transcriptional mechanisms is essential. We present here an efficient implementation for analyzing co-transcriptional RNA-ligand interaction dynamics. This approach allows for the first time to model concentration-dependent metabolite binding/unbinding kinetics. We exemplify this novel approach by means of the recently studied I-A 2’-deoxyguanosine (2’dG)-sensing riboswitch from *Mesoplasma florum.*

## 1. Background

Riboswitches are cis-acting regulatory RNAs that undergo an allosteric conformational switch upon binding of a cognate metabolite. They have originally been characterized in bacteria, where they are typically located in 5’ untranslated regions (5’-UTR) of mRNAs. The regulatory repertoire of procaryotic riboswitches comprises modulation of the expression of adjacent genes by means of transcription termination or translation initiation. Riboswitches typically consist of two domains, an evolutionary conserved aptamer domain that specifically senses a metabolite and a variable expression platform that can form specific RNA structural elements required for modulation of gene expression [26, 13]. The binary characteristics of a transcriptional ribsowitch, being either premature transcription termination (OFF-switch) or continuation (ON-switch) is triggered by metabolite binding. In this line, aptamer formation, ligand binding and subsequent formation of RNA structures in the expression platform (terminator / anti-terminator) are kinetically controlled co-transcriptional events [32].

Conceptually, a ligand-binding domain senses the concentration of specific small molecule metabolites and translates this information into a structural change of the 5’-UTR region, which in turn modulates mRNA expression. The engineering of RNA based regulation and the rational design of complex RNA regulatory circuits is an attractive challenge from a synthetic biology point of view, because both directed-evolution based experimental techniques and efficient computational approaches are well developed.

The model for co-transcriptional RNA folding outlined here is based on several assumptions that allow for a reasonable balance between accuracy and computational feasibility. We describe RNA at the level of secondary structures, since the majority of the free energy of RNA folding is captured by secondary structure formation. This level of description does not explicitly account for tertiary interactions, nor can RNA-ligand binding be modeled from first principles. Instead we rely on experimentally determined binding motifs and affinities. Nevertheless, this provides a computationally tractable yet appropriate approach for modeling equilibrium and non-equilibrium properties of RNA [23, 37]. Due to the system size and time-scales of the transcription process, a fully atomistic treatment using molecular dynamics is currently out of reach. While there is much recent interest in coarse grained methods for RNA 3D modeling [29], these approaches still lack necessary reliability and accuracy.

Co-transcriptional RNA structure formation has long been recognized as a core process mediating RNA function [1, 19, 25]. Nevertheless, accessible computational approaches for reliably modeling the folding behavior of nascent RNA are still scarce. While early approaches such as RNAkinetics [2] or Kinefold [35] are available as web services, they do not provide sufficient flexibility for ligand support [5] and cannot be modified due to lack of available source code. Zhao et al. [36] have proposed a helix-based approach for computational modeling of co-transcriptional RNA folding. They derive a set of structures by sampling and estimate rates between these states from enthalpic and entropic contributions of helix formation and disruption. Limiting elementary moves to entire helix operations dramatically reduces the conformation space. In a recent paper they added ligand binding support to their model [10], however they do not provide software. In this contribution, we therefore rely on in house developed software, based on the `ViennaRNA` package [23].

Recently, two different chemical probing techniques have been established, that make co-transcriptional folding more accessible to experiment. Watters et al. [30] employed SHAPE probing of co-transcriptional intermediates, while Incarnato et al. [17] exploited DMS chemistry. Both achieved near-nucleotide resolution of co-transcriptional folding intermediates, however, time-resolution is still limited, so that only longer lived intermediates can be detected.

## 2. Motivation

The computational design and analysis of riboswitches focuses mainly on thermodynamic aspects of the full length regulatory element. These approaches ensure that the designed RNA element is optimal with respect to a cost function specifying multiple sequence, structure and energetic constraints [14, 3]. However, RNA folding starts already during the transcription process under normal cellular conditions. The superposition of two dynamic processes, i.e. chain-growth and kinetic folding of the already transcribed chain piece makes the overall process history dependent and has therefore a huge impact on which structure is adopted by the fully transcribed RNA chain. Moreover, in riboswitches it is the interaction with the ligand that determines the correct final structure. This complicates the description, as RNA-ligand interactions are not part of the standard RNA secondary structure model.

In this contribution we present an efficient approach for prediction of co-transcriptional RNA folding in the presence of a ligand. Ligand binding is modeled as an all-or-none process, i.e. a ligand can only be bound by a fully formed aptamer structure. In this line, a structure is considered binding competent if and only if it contains the complete aptamer. This neglects ligand-binding by partially formed aptamer intermediates, for which affinities are not available anyway. We use experimentally determined binding affinities rather than modeling them from first principles.

The rationale for this type of modeling is learning design rules for de-novo or re-design of riboswitches in synthetic biology applications. We therefore selected the type I-A 2-deoxyguanosine (2dG)-sensing riboswitch from *Mesoplasma florum,* which has recently been studied by NMR spectroscopy [15], as an example here and demonstrate that our results are in good agreement with experimental data. Computational predictions yield significant additional mechanistic insight, since the NMR experiment only captures the equilibrium structures for a number of transcriptional intermediates rather than the full co-transcriptional dynamics at the level of single nucleotide chain extension events. This allows not only to understand natural riboswitches on mechanistic level, but also perform systematic *in silico* screening and analysis of novel designs [3] prior to expensive experimental validation.

## 3. RNA folding dynamics

### 3.1 General Concepts

RNA folding can be seen as a dynamic process on an energy landscape, consisting of a set of conformations (here secondary structures), each of which is assigned a (free) energy value. In addition, a move set defines which structures are connected by elementary transitions. In the following we will assume the simplest move set allowing only for opening or closing of single base pairs. Each of these transitions has an associated rate k. We will assume a simple Metropolis rule for these rates, such that a transition from some conformation x to a neighbor y is given by

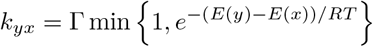

The prefactor Γ sets the time scale of the process and can be estimated by comparing with experimentally determined refolding rates. Comparison to NMR measurements of a small bistable RNA [7] suggest Γ ≈ 10^−5^*s*^−1^.

### 3.2 Kinetic folding of RNA

Folding dynamics can then be modeled by a continuous-time Markov process based on a master equation which describes the change in state probabilities *P_t_*(*x*) to see state *x* at time *t*

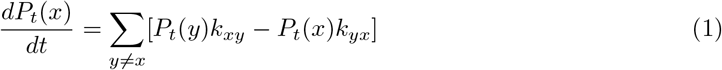

The system of linear differential equations can be re-written in matrix form 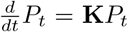, with the formal solution

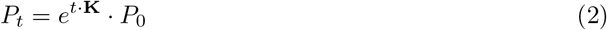

Solving eq. 1 directly is impractical for anything except toy examples, since the dimension of the rate matrix **K** is equal to the number of structures in the conformation space and computing the matrix exponential eq. 2 therefore becomes unfeasible. One possibility to address this issues is to approximate the solution of the master equation through stochastic simulations using a Monte Carlo method. This approach is taken e.g. by `kinfold` [4]. While the outcome of this method can be regarded as gold standard, computing and analyzing a large number of trajectories can be time consuming and tedious. An alternative approach is to reduce the number of states (and thus the dimension of the matrix **K**) through coarse graining, as done by the `barriers` program [6]. The treekin program [33] provides an efficient solver for eq. 2 via matrix diagonalization, that works well for up to 10000 states.

Another important, yet often neglected, aspect is the fact that RNA structure is formed already during its synthesis, i.e. it folds back on itself co-transcriptionally. Co-transcriptional folding is fairly easy to implement in simulation approaches [12, 4]. In the landscape view, co-transcriptional folding induces a landscape that varies over time. The `BarMap` framework explicitly models such a scenario [16]. Finally, methods such as `kinwalker` [8] attempt to construct a single, most likely, folding trajectory for the growing RNA chain. While this introduces fairly drastic approximations it can be applied to RNAs up to ≈ 1500 nt length.

### 3.3 The kinfold approach

The dynamics of RNA folding can be modeled by a Markov process using the opening and closing of single base pairs as the neighborhood function to change state. The program `kinfold` [4], available via the `ViennaRNA` package [23], provides an efficient Gillespie-type simulator for the stochastic folding of RNA molecules. A simple extension allows to simulate co-transcriptional folding of the growing chain by appending the next nucleotide of the sequence at regular time intervals until the full length is reached. As with all direct simulation approaches, a large number of trajectories has to be collected and analyzed in order to get statistically significant results. A typical way to analyze the trajectories is to define structural classes, characterized by the presence or absence of structural features of interest (such as binding motifs or terminator hairpins). The sampled trajectories can then be projected into these structural classes ( see Figure 1), to obtain summary statistics on the frequency of features over time. An example of such an analysis is shown in Figure 2.

**Figure 1:**
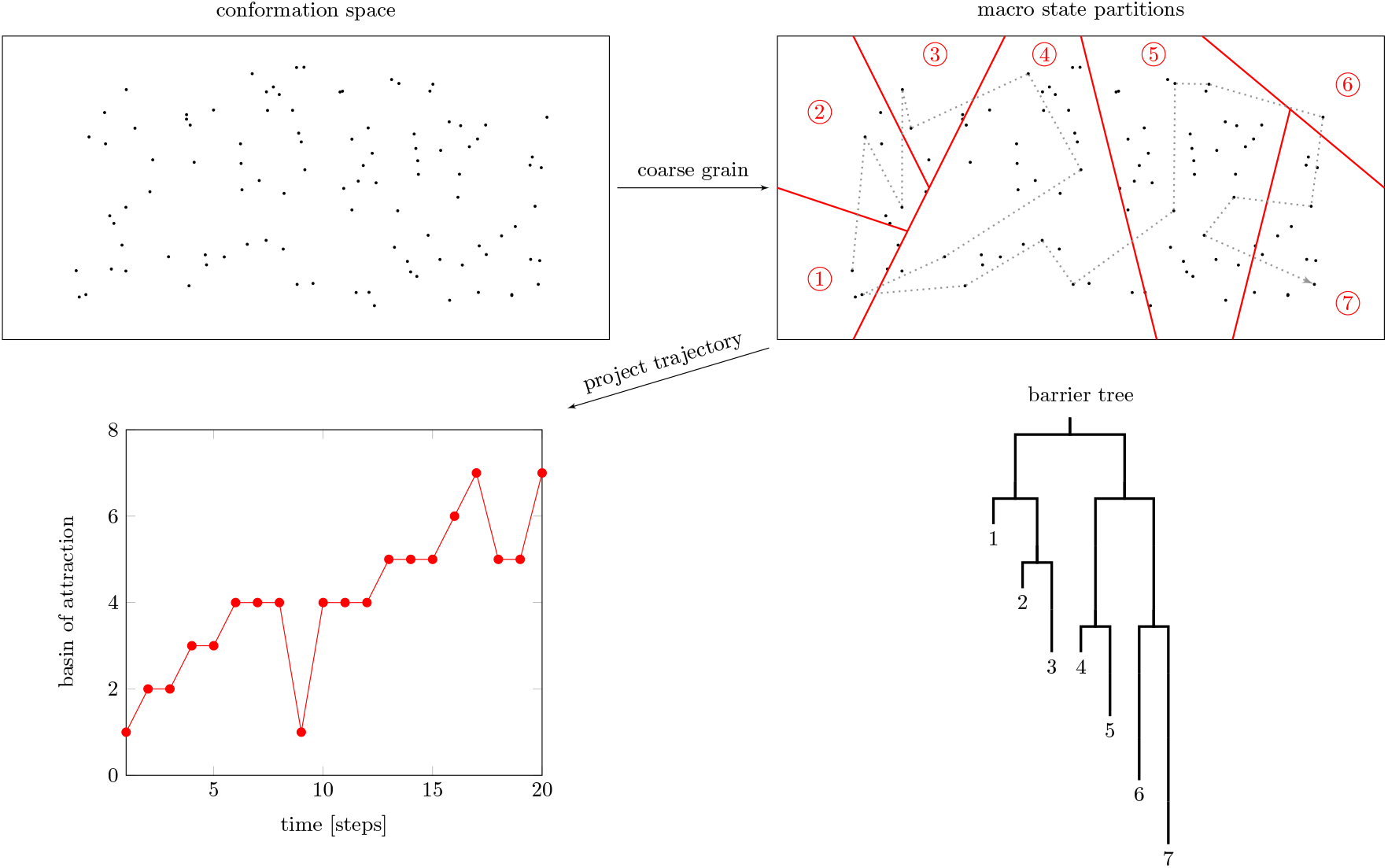
RNA folding landscape abstractions. The different secondary structures an RNA sequence can fold into form a discrete energy landscape, where insertion/deletion of single base pairs induces a valid neighborhood relation between states in the landscape. The process of RNA folding can be described as a Markovian walk between the states in this conformation space (dotted line). The full conformation space is, however, too large to study the folding dynamics of an RNA molecule. Therefore, a unique partitioning of the states into much fewer basins of attraction (areas marked with circled red numbers) is performed. The folding dynamics can then be described in terms of transitions between these macro states. The macro state partitioning can be visualized in the form of a barrier tree, where leaf nodes (basins of attraction) are connected by the energetically lowest saddles points.

**Figure 2:**
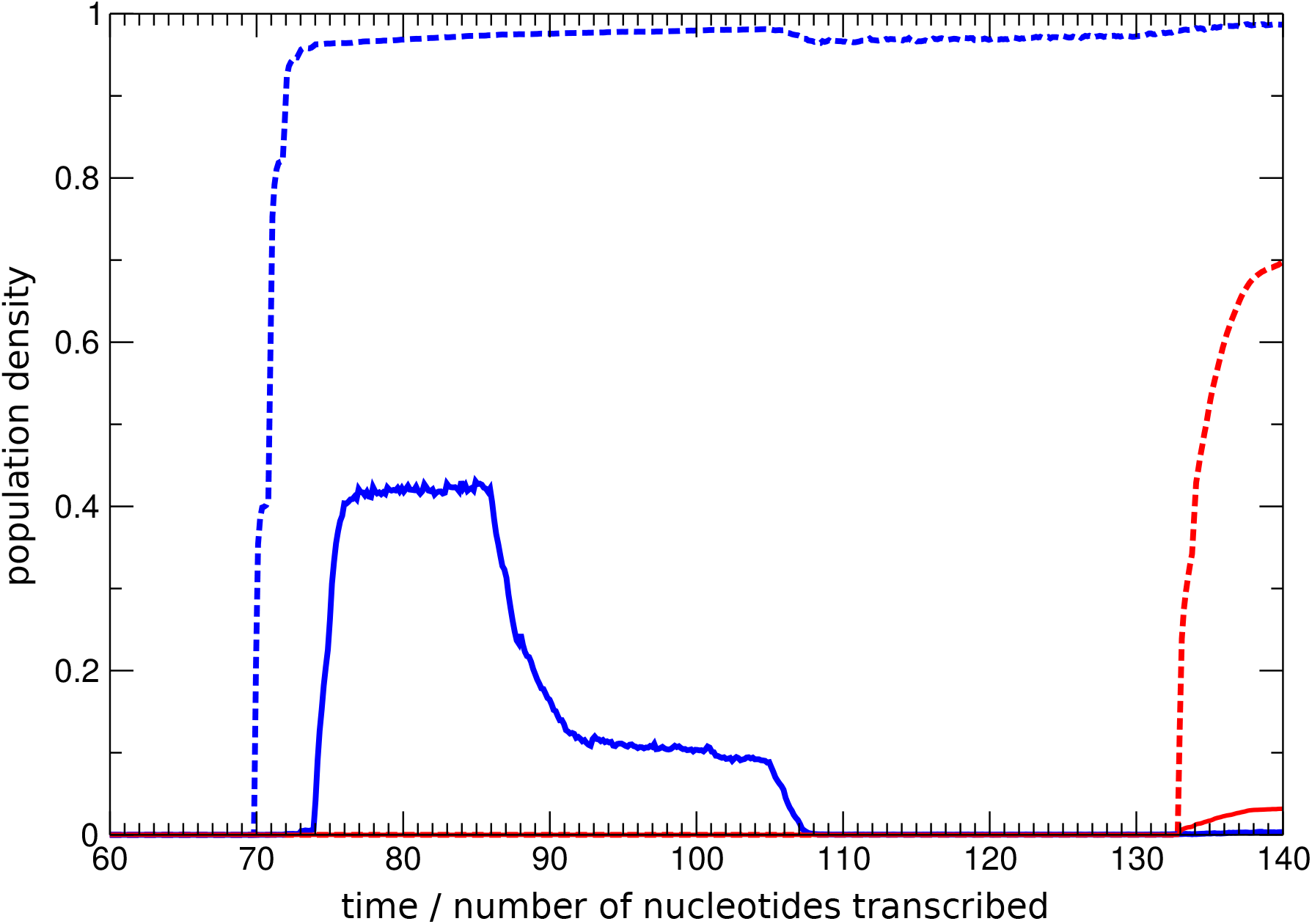
Summary statistics of kinfold simulations of the dG riboswitch. Dashed/solid lines indicate simulations with/without ligand present. The plot illustrates the aggregated probability density of aptamer (blue) and terminator (red) motifs present in the ensemble of folding RNA molecules *versus* time measured in numbers of nucleotides transcribed. If the ligand is present, the aptamer motif shows a quick increase in density to a value of almost 1, which remains stable upon further chain elongation.

### 3.4 The barriers approach

While `kinfold` simulations are the most fine grained approach, generating a statistical representative ensemble of trajectories can be quite time and resource consuming. Another drawback of the `kinfold` approach is that the partitioning of the structures along trajectories to calculate the time dependence probability to find a particular structural feature in the ensemble has to be done manually. The program `barriers` [6], in contrast, performs a partitioning of structures into feature classes automatically. In detail, a unique coarse-graining of the conformation space is constructed by assigning each structure *x* to the local minimum that forms the end-point of a gradient walk on the energy landscape starting from *x*. The resulting macro-states can be naturally represented by a tree, with leafs corresponding to local minima, connected by internal nodes representing the energetically lowest saddle point between them, see Figure 1. In addition, `barriers` computes effective rates between the macro states, which can be used to solve the master eq. 1 using `treekin` [33].

Since `barriers` relies on an exhaustive enumeration of low energy structures, the computational effort grows exponentially with sequence size, which limits the length of RNAs that can be studied to a little over 100 nt. Recently, a number of heuristic approaches have been reported that attempt to raise this limit based on flooding techniques [34, 24] or sampling of local minima [27, 28, 20, 21].

The `barriers` program analyses a single landscape of fixed length. Co-transcriptional folding, however, can be seen as a process that happens on a sequence of landscapes of different length. This is exploited by the `BarMap` pipeline [16]. In short, `BarMap` defines a mapping between macrostates of successive landscapes. This mapping is used to transfer the final population density from a `treekin` simulation on the landscape at length l to the following landscape (with length l + 1), yielding the starting condition for the folding simulation on the extended landscape. The time dependent population densities from each simulation can then be concatenated to obtain the full co-transcriptional folding profile, see Figure 4.

### 3.5 RNA-Ligand interaction dynamics

RNA-ligand interactions are crucial for the functioning of cis-acting ribo-regulators, but are outside the conventional RNA secondary structure model. Moreover, while RNA (re-)folding can be seen as isomerization, ligand binding is a bi-molecular reaction and therefore depends on the concentrations of both RNA and ligand. Nevertheless, binding of small molecules to aptamers can be included in our models, if the structural prerequisites for ligand binding (in form of a secondary structure motif), as well as the free energy of binding are known from experiment. In the simplest case, the energy model is adjusted, by adding the binding energy as a bonus to all binding competent structures. Such an approach is taken in the `kinfold` simulations shown in Figure 2. This neglects all concentration dependencies and also assumes that ligand binding is instantaneous (i.e. much faster than opening or closing of base pairs).

A more realistic scenario treats RNA-ligand binding as a separate reaction channel. Here, the case where ligand is present in large excess is particularly easy to handle, since the concentration of free ligands is then constant over time. Kuhnl et al. [22] worked out the theory how to treat this case in the `barriers` framework. However, they only provide a proof-of-concept implementation that runs `barriers` on both the ligand-free and fully-bound landscape, dumps all macro-states to disk and re-reads them in a Perl script that computes cross-rates between bound and unbound macro-states. The dG riboswitch example presented here would be too large to compute with their implementation. For this work we therefore provide an efficient implementation of the Kühnl approach as new version of the `barriers` program.^1^

We assume that the binding free energy *θ* as well as the dissociation rate *k*_off_ = *k*_on_*e*^−*θ/RT*^ are known from experiment. The rate of complex formation becomes proportional to the ligand concentration *C, r* = *k*_on_ ·*C*. We construct the input for the `barriers` program such that it contains ligand-bound and unbound versions of each binding competent structure. `barriers` then constructs the macro-states separately for the bound and unbound components of the landscape, as well as the transition rates within each component. Finally we compute, the transitions between bound and unbound states assuming a value of 1 for both *k*_on_ and *k*_off_ (i.e. ignoring all concentration or binding energy effects). We thus obtain a transition matrix consisting of four distinct blocks: Two blocks corresponding to transitions within bound / unbound states and two blocks for binding and unbinding transitions (black, blue, red and green blocks in Figure 3). The binding and unbinding rates can now be adjusted in the transition matrix for a particular ligand concentration, as well as the values for *k*_on_ and *θ* in a simple post-processing step. This has the advantage that the computation can be repeated for many different concentration values, without the expensive step of re-computing the macro-state partitioning.

**Figure 3:**
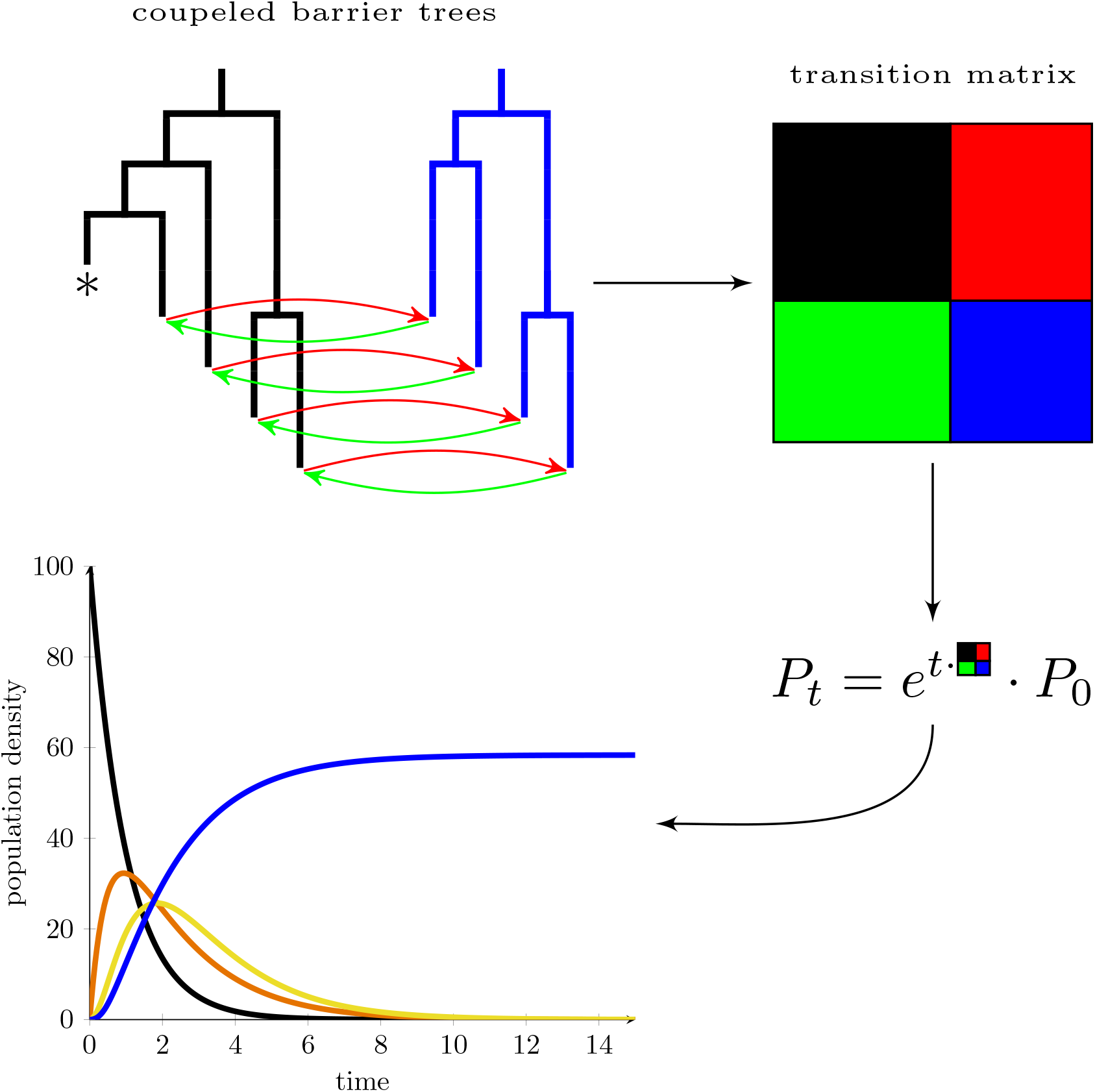
Simulation of the folding dynamics. The program `barriers` partitions the folding landscape into two classes of structures: (i) the aptamer domain is present and the ligand is bound (blue barrier tree) and (ii) the ligand is not bound (black barrier tree). The latter class contains also structure that do not form the aptamer binding domain (marked with *). Binding/unbinding events of the ligand to binding competent structures (red/green arrows) factors out to transitions between the corresponding leaf nodes of the two barrier trees. The transition matrix of the whole system, therefore, decomposes into distinct blocks. Transitions within the trees are located along the diagonal (black/blue areas) while transitions between trees are found in outer-diagonal regions (red/green areas). Substituting the transition matrix into the formal solution of the master equation 2 and varying the time t yields the information how the population density of basins of attraction changes over time.

## 4. Results

### 4.1 Showcase: dG riboswitch

Purin-binding riboswitches (Rfam family RF00167) form an abundant class of regulatory elements in bacteria, that can sense guanine, adenine, or 2’dG. Here, we use the type I-A 2-deoxyguanosine (2dG)-sensing riboswitch from *Mesoplasma florum* (mfl-riboswitch) as showcase for a transcriptional OFF-switch under kinetic control [18]. The system discussed here regulates expression of a downstream ribonucleotide reductase in *M. florum* and is yet one of two known representatives of dG-sensing riboswitches. The other one has recently been reported in *Gracillimonas tropica* [31].

In a transcriptional OFF-switch, a terminator hairpin leads to pre-mature transcription termination, but only in the presence of the ligand. Folding the 143 nt leader RNA sequence with RNAfold [23], however, shows that the terminator stem is present in equilibrium even without additional ligand stabilization. This raises the question how this riboswitch can ever be in an ON state. Apparently, the regulatory decision has to be taken before the RNA polymerase reaches the termination site, indicating that the switch is under kinetic control.

In a recent study, the regulatory potential of the *mfl*-riboswitch has been investigated in detail by NMR spectroscopy [15]. Equilibrium NMR structures of different length intermediates coincide with computationally predicted equilibrium states. As shown in the study, a transcript of the first 75 nt forms a stable aptamer structure. At a length of around 110 nt. however, the aptamer structure is no longer stable and instead an alternative (anti-terminator) structure is formed. At 133 nt a structure containing both the aptamer and the terminator hairpin becomes the most stable conformation. While the experimental analysis of equilibrium structures alone cannot explain the riboswitch function, it suggests a scenario, where without ligand anti-terminator containing structures are formed at around 110 nt, and where subsequent terminator formation is too slow to disengage the polymerase. In the presence of ligand, the aptamer structure is strongly stabilized, thereby inhibiting formation of the anti-terminator and allowing fast formation of the terminator starting at length 133. Without an experiment that directly probes folding *dynamics,* only computational simulation can be used to corroborate this scenario.

### 4.2 Co-transcriptional folding dynamics with kinfold

We first studied the co-transcriptional folding of the dG ribsowitch using `kinfold`. We used the simplest model for treating ligand binding, by defining an aptamer motif and assigning a bonus energy of 8kcal/mol to all structures containing the motif. This corresponds to assuming an infinite ligand concentration as well as infinitely fast binding. The value of 8kcal/mol was chosen as reasonable approximation, since experimental data for 2’-deoxyguanosine binding are not available [9].

We simulated two ensembles (with and without ligand) of 10000 trajectories each using `kinfold`. Each structure along the trajectory was then classified according to the presence/absence of the aptamer motif and presence/absence of the terminator hairpin, thus defining four possible structural classes. This data was then used to calculate for every time point the probability to find the folding chain in an aptamer or in a terminator containing state (blue and red lines in Figure 2). In the absence of ligand (solid lines) the aptamer motif (blue line) starts forming after 70 nucleotides have been transcribed, and reaches a maximum density of 0.4 around 85 nucleotides. Upon further chain growth the aptamer motif becomes unstable and vanishes around 105 nucleotides from the population of folding molecules. The terminator motif (red line) very slowly gains population density starting around length 133. The folding behavior changes dramatically in the presents of the ligand (dashed line). In this case the aptamer motif forms around time 70 and very quickly rises to a density of almost 1, stably staying at this level until all nucleotides have been transcribed. Around 135 transcribed nucleotides the terminator motif shows a sharp increase in density, presumably leading to effective transcription termination.

### 4.3 Co-transcriptional interaction dynamics with BarMap

In the next step, we added ligand support to the `BarMap` [16] pipeline^2^ to simulate the co-transcriptional interaction dynamics of the dG riboswitch. As described above this allows us to explore the effect of ligand concentration on switching behavior.

Without ligand, the barmap simulations showed behavior qualitatively similar to the `kinfold` simulations above (data not shown). The population is dominated by aptamer containing conformations at lengths around 80. At 90 nt the density of aptamer conformations drops sharply and reaches 0 at 110 nt. The lost density is shifted into anti-terminator containing conformations, which remain dominant until 140 nt, thus delaying formation of the terminator. At very high ligand concentrations, the behavior is consistent with the `kinfold` simulation with ligand (dashed lines in Fig. 2). The aptamer is populated at almost 1 and remains until the full chain is transcribed.

A more realistic scenario, corresponding to a physiological ligand concentration of 1 mM, and a binding energy of 8 kcal/mol is shown in Figure 4. Here, the co-transcriptional dynamics of the dG riboswitch is dominated by three mutually exclusive folding intermediates: aptamer domain, anti-terminator and terminator states, which are represented in shades of blue, green and red, respectively. A “pre-aptamer” structure at length 74 is populated to more than 0.5. Since the closing hairpin of the aptamer is not yet fully formed at this chain length, this structure is not considered binding competent in our simulation (dashed lines). Addition of one nucleotide renders the aptamer structure complete, making available ligand-bound states. In our simulation, population density is shifted to metabolite-bound aptamer structures at length 75. While the aptamer conformation is lost around length 110, unbound intermediate conformations with structural properties of the anti-terminator dominate the ensemble between lengths 110 and approx. 133 (dashed green lines). Subsequent chain elongation past nucleotide 133 shifts the population density towards terminator structures, corresponding to the thermodynamic ground state (red lines).

**Figure 4:**
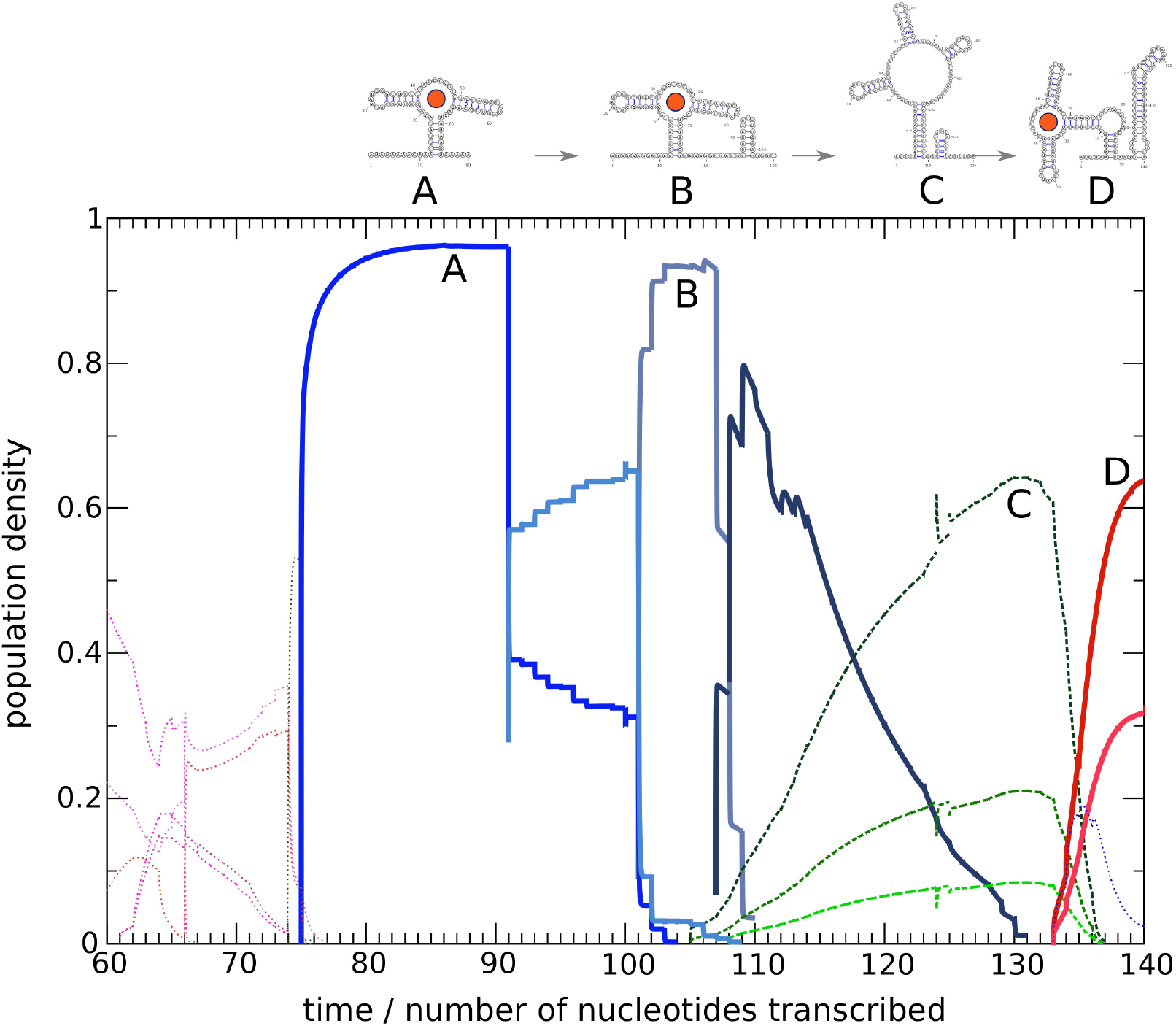
Co-transcriptional interaction dynamics of the dG riboswitch. A `BarMap` simulation with explicit ligand concentration of 1 mM and stabilization bonus of 8 kcal/mol was computed at 298 K for chain lengths 20 — 143 (only the relevant region from lengths 60nt-140nt and macro states with population above 0.05 are shown). Ligand-bound states are plotted bold, unbound states are dashed. Blue lines starting from chain length 75 represent states that form the aptamer, while green lines represent anti-terminator-like states. Refolding to the terminator-dominated ensemble (red lines) starts from chain length 133. Selected intermediate secondary structure representatives are plotted above, associated to macro state population curves by capital letters A-D.

## 5. Materials and Methods

All computer simulations of the co-transcriptional folding of the dG riboswitch were performed at 298K (in accordance with NMR experiments from a previous study [15]) based on the `ViennaRNA` package v2.4.3 [23].

### 5.1 kinfold Monte Carlo simulations

`kinfold` (shipped with the `ViennaRNA` package) was used to simulate 10000 trajectories of the growing dG riboswitch with/without ligand starting from an initial chain length of 40 nt. A bonus energy of 8 kcal/mol was added for ligand stabilization. Comparison with experimental data [7] suggested that 1 s corresponds to approx 100000 internal time units. Hence a chain elongation event every 4000 simulation steps is equivalent to a transcription rate of 25 nt/sec.

### 5.2 BarMap coarse grained simulations

Coarse-grained co-transcriptional folding dynamics simulations were performed with `BarMap` [16]. The `BarMap` pipeline consists of multiple computational blocks that build on one another, i.e. landscape generation, macro state mapping and kinetics simulation by density remapping. Ligand support in the landscape generation part is achieved by classifying binding competent structures in the conformation space for each growing chain fragment starting from chain length 75, which corresponds to the fully formed aptamer. We required the secondary structure motif of the full aptamer to be present at the 5’ end in order to qualify binding competent structures. Initially, the lower part of the energy landscape was exhaustively enumerated for each growing chain intermediate with `RNAsubopt` up to 17kcal/mol above the ground state. Structures with lonely base pairs were excluded from our simulation in order to reduce the size of the conformation space.

We used `barriers` v1.7.0 to perform the macro state decomposition and for computing block-diagonal transition matrices. Construction of the energy landscape was performed with a minimum barrier height of 1.5kcal/mol, thereby considering up to 1500 macro states. If the resulting energy landscape components (i.e. ligand-bound and unbound) were not connected among themselves, the energy range for conformation space enumeration was increased by 1 kcal/mol until the landscapes were connected.

Given that we disregard structures with isolated base pairs results in the fact that both single moves (i.e. insertion/deletion of a single base pair) and double moves (i.e. insertion/deletion of a double base pair) are possible in order to inter-convert one RNA structure into another. This, however, has to be taken care of when computing (microscopic) transition rates within `barriers`. A double move represents two insertion/deletion events, hence the corresponding rate should be slower than that of a single move. We therefore scaled all double moves with a factor 0.01 to account for this inconsistency towards faster kinetics of double moves.

While `barriers` does not allow for transitions between bound and unbound states during macro state construction, we implemented a Perl script that does this computation in a post-processing step. `crossrates.pl` is shipped with `barriers` v1.7.0 and we applied it to account for different ligand concentrations and stabilization energies. For the data shown in Figure 4, physiological conditions in the form of 1 mM ligand concentration and 8kcal/mol stabilization energy was chosen. Following [22], we used the default association/dissociation constant *k*_on_ = 600 M^−1^s^−1^.

Starting from an initial sequence length of 20 nt, we computed the population dynamics for each growing chain element for 4000 time steps until the chain was fully elongated. The choice of a maximum of 1500 macro states for each energy landscape results in a very fine-grained resolution of the individual folding dynamics runs. However, this leads to many macro states with low population density, which makes it difficult to interpret results. To overcome this problem, we applied a coarse-graining strategy as a post-processing step. The simplest criterion for such a coarse graining is discrimination by barrier height, i.e. all macro states that are separated by an energy barrier below a certain threshold are merged for better visualization of the `BarMap` results. Here, we used 3kcal/mol as minimum barrier height.

The time and space limiting step of the procedure remains the analysis of individual length landscapes using `barriers` [6]. The latter processes an exhaustive list of low energy secondary structures, here within a range of 18 kcal/mol above the ground state. For each input structure `barriers` constructs the set of neighbors that can be reached by a single move (opening / closing of a single base pair), the computational effort as well as the memory requirements therefore scale as 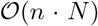, where *n* is the sequence length and *N* the number of input structures. For the dG riboswitch analyzed here, the largest landscape comprised 2.04 billion structures and took 98 hours to process. Since the landscape size grows exponentially with sequence length, the 143 nt dG riboswitch is close to the limit of what can be analyzed on typical current hardware.

## 6. Discussion

We have presented a novel approach for simulating co-transcriptional folding in the presence of ligands. Support for ligand binding is essential for understanding the mechanism of riboswitches, but is unfornately unavailable in most existing tools for RNA folding kinetics. The approach extends our `barriers-BarMap-treekin` pipeline and is implemented as freely available software.

Most recently, the Zhang group has applied their kinetic folding model also to co-transcriptional folding with ligands [10, 11]. This is a much more coarse grained, helix based model in which the landscape is presented by just a handful of conformations. It is thus unclear, whether all relevant states are included. Nevertheless, they present promising results on several example riboswitches. In contrast, our method starts from an exhaustive set of low energy structures and is therefore free of *a priori* assumptions about relevant states or folding pathways. Instead, all relevant metastable states emerge as a result of the computation. Only the binding competetent sub-structure of the aptamer has to be known in advance.

Current approaches for *in silico* design of RNA switches focus on thermodynamics design, but do not take co-transcriptional folding dynamics into account. However, as exemplified by the dG riboswitch above, the mechanism of transcriptional switches cannot be understood from thermodynamics alone. The superposition of re-folding and chain elongation processes makes co-transcriptional folding inherently a history dependent process.

Directly incorporating these approaches into the objective function of a sequence design procedure may still be computationally challenging. At the very least they can, however, be used to filter sequences suggested by purely thermodynamic design methods. In this line, given that it is difficult to explain co-transcriptional folding or re-folding mechanism with experimental methods alone, our approach can be used to assess the ligand-dependent dynamics of designed sequences before they are tested in the wet lab.

A simple thermodynamic design of an OFF-switch similar to the dG riboswitch would use the following criteria: (i) at an early length *l*_apt_, aptamer containing structures dominate the equilibrium ensemble, *p*(apt) ≈ 1 (ii) at intermediate lengths the aptamer should vanish, *p*(apt) ≈ 1 (iii) at full length structures containing both aptamer and terminator should be dominant, *p*(apt) ≈ 1 Λ *p*(term) ≈ 1. These criteria can easily be expressed in an objective function for RNAblueprint [14].

Co-transcriptional folding simulations suggest that additional kinetic criteria are necessary: (iv) binding has to be sufficiently fast (tunable by the ligand concentration), such that aptamer structures are bound and stabilized before they are lost at intermediate lengths and (v) unbinding (tunable by the binding energy *θ*) has to be sufficiently slow, such that a ligand bound aptamer survives for many chain elongation steps, even if the unbound aptamer is no longer stable.

In terms of rates, these criteria translate into the inequalities

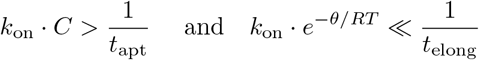

where *t*_apt_ is the time window during which the aptamer containing structures dominate the equilibrium ensemble, and *t*_elong_ is the time for one chain elongation step.

A remaining limitation for switch design is the relatively small number of aptamers for which binding energies, let alone binding rates, have been determined experimentally.

## 7. Conclusion

We have added support for ligand binding/unbinding dynamics to the kinfold [4] and BarMap [16] tools and have demonstrated novel functionality of our software by means of a relatively challenging riboswitch example, which has recently been studied by NMR spectroscopy [15]. Our results suggest that we can reproduce and explain the experimentally determined switching behavior of the dG riboswitch. More generally, if binding affinities and the binding competent aptamer structure of a riboswitch are known, our method can be used to model the co-transcriptional folding dynamics with and without ligand. A future perspective of this approach is inclusion of co-transcriptional time profiles, that resemble transcriptional pausing often observed in nature.

All software presented in this contribution is publicly available through the ViennaRNA channel on Github at https://github.com/ViennaRNA.

## Acknowledgments

This work was partly funded by the Austrian science fund FWF project F43 “RNA regulation of the transcriptome”.

## Footnotes

[1] version 1.7, https://github.com/ViennaRNA/Barriers

[2] https://github.com/ViennaRNA/BarMap

